# Stress-responsive enhancer RNAs couple chromatin reprogramming to post-transcriptional control of senescence

**DOI:** 10.64898/2026.03.20.713302

**Authors:** Rene Kuklinkova, Natalia Benova, Jaskaren Kohli, James R Boyne, Wayne Roberts, Chinedu A. Anene

**Author notes:** Corresponding author. Centre for Biomedical Science Research, School of Health, Leeds Beckett University, Leeds LS1 3HE, United Kingdom. These authors contributed equally (Rene Kuklinkova, Natalia Benova).

## Abstract

**Background:** Cellular senescence, a driver of ageing and age-associated disease, is accompanied by extensive epigenomic reprogramming that changes enhancer RNA levels, yet how enhancer activity is translated into functional RNA-level regulation remains unclear. Here we investigate how this enhancer reprogramming impacts functional RNA-level regulation by eRNAs.

**Results:** By integrating time-resolved transcriptomic analyses across multiple primary human cell types, we identify recurrently dysregulated senescence-associated enhancer RNAs (SAeRs). We focus on EN526, which is reproducibly repressed during senescence while its locus remains broadly stable across cell states. EN526 exhibits cytoplasmic localisation and extensive eRNA–mRNA interactions, and cytoplasmic depletion of EN526 recapitulates its senescence-associated loss and alters the stability and translation of the cell-cycle regulator CDKN2C. EN526 perturbation further mediates stress responses, cellular survival, and extracellular remodelling associated with the senescence phenotype.

**Conclusion:** Together, these findings show that SAeR changes accompanying enhancer reprogramming in senescence are not merely passive events but can act as functional intermediates linking enhancer dynamics to post-transcriptional regulatory networks that phenocopy key senescence-associated cellular features. Extending this model, genetic associations at the EN526 locus further connect this regulatory axis to age-related traits and circulating protein phenotypes, supporting its broader relevance to human ageing and disease.

## 1. Introduction

Cellular senescence is a complex and multifaceted stress response that contributes to tissue homeostasis, development, and tumour suppression, while also driving age-associated dysfunction and chronic disease ^1–3^. Although senescence arises from diverse triggers, including replicative exhaustion, oncogenic signalling, and genotoxic stress, its establishment is consistently accompanied by widespread transcriptional and chromatin reorganisation ^4–6^. Senescent cells undergo large-scale nuclear remodelling, heterochromatin redistribution, and enhancer reprogramming ^7–9^, which together reshape gene expression programmes that stabilise cell-cycle arrest, inflammatory signalling, and altered tissue behaviour ^6,10^. While these chromatin-level changes are well documented, how they propagate into downstream regulatory layers that sustain the senescent phenotype remains incompletely understood.

Enhancers are central integrators of developmental, stress, and inflammatory signals ^11,12^, and their activation state is extensively remodelled during senescence ^13–16^. Transcription from these elements produces enhancer RNAs (eRNAs), a class of non-coding transcripts increasingly recognised as regulators of transcriptional and post-transcriptional processes. In some contexts, enhancer transcription itself can influence chromatin accessibility and gene activation ^17^, whereas in others the RNA molecules contribute directly to regulation, for example by stabilising enhancer–promoter interactions, recruiting transcriptional regulators, or modulating cofactor availability ^18–21^. These observations suggest that eRNAs may act as molecular conduits through which enhancer reprogramming shapes senescent phenotype.

Emerging evidence further indicates that eRNAs can function beyond transcriptional control. Several cell fractionation datasets detect cytoplasmic localisation of subsets of eRNAs ^22,23^, and recent works, including our own, demonstrates that such transcripts can engage in eRNA–mRNA interactions that influence mRNA stability and translation independent of transcription ^24^. These findings expand enhancer function into the post-transcriptional space, suggesting senescence-associated enhancer reconfiguration could influence cellular phenotypes through non-coding RNA-mediated mechanisms as well as chromatin-level control. Despite these advances, the contribution of eRNAs to the senescence phenotype remains poorly defined. Previous studies have linked enhancer activity to senescence-associated transcriptional programmes, yet whether eRNAs themselves participate in shaping senescence, particularly through RNA-level regulatory mechanisms, is unknown. Given that senescence involves coordinated rewiring of chromatin architecture, transcription, and RNA processing, eRNAs represent strong candidates for mediating communication between these layers.

Here we investigate how enhancer reprogramming during senescence translates into functional RNA-level regulation at the post-transcriptional level. By integrating time-course transcriptomic analyses across multiple senescence cell types, we identify a set of recurrently altered senescence-associated eRNAs (SAeRs). We found that one of these transcripts (EN526) exhibits cytoplasmic localisation and post-transcriptional regulatory activity that phenocopies senescence features, and we define repression of EN526-CDKN2C axis, as a central node linking enhancer regulation to eRNA control of senescence fate through altered CDKN2C, PLAU and COL18A1. Together, this work positions eRNA regulation as active components of the senescence regulatory network and reveals a molecular pathway connecting enhancer reconfiguration to post-transcriptional control through eRNAs.

## 2. Methods

### 2.1 Integrative analysis of omics datasets across senescence contexts

We integrated a range of publicly available transcriptomic, epigenomic, and proteomic datasets to investigate eRNA regulation and downstream molecular effects during cellular senescence. RNA-sequencing datasets representing irradiation-induced senescence across primary human fibroblasts, keratinocytes (NHEK), and melanocytes were analysed to identify recurrently dysregulated eRNAs and define SAeRs (Fig. 1a–c; ENA: E-MTAB-5403; poly(A)-enriched ^6^). An independent transcriptomic dataset spanning proliferating, quiescent, senescent, deep senescent, and senescence-reversal states in BJ fibroblasts was analysed to validate EN526 repression across senescence conditions (Fig. 1d; GEO: GSE133292; poly(A)-enriched ^25^). Chromatin accessibility and histone modification profiles at the EN526 locus were examined using ATAC-Seq and ChIP-Seq datasets from the same experimental model (Fig. 1e; GEO: GSE133292). To investigate transcriptional regulation of EN526, ChIP-Seq binding profiles from ReMap were analysed to evaluate evidence for TF occupancy at the EN526 locus (Fig. 1g–h; ReMap2022 ^26^). Transcription factor perturbation followed by RNA-Seq from ENCODE and GEO were analysed to evaluate changes in EN526 expression following knockdown of BACH1 and SETBP1 in K562 cells, knockdown of HMGA1 in KYSE-30 cells and overexpression of JUNB in A375 cells (Fig. 1i; Encode: ENCSR226CVP and ENCSR109KMO ^27^, GEO: GSE159539 and GSE263439 ^28,29^). RNA-Seq datasets from IMR90 fibroblasts spanning proliferating, quiescent, and senescent cells induced by etoposide or irradiation were further analysed to examine post-transcriptional and extracellular outputs of the EN526-CDKN2C axis (Fig. 2e-f, 4c-e; GEO: GSE227766 ^30^). Ribosome profiling data from the same experimental model were used to assess translation efficiency of CDKN2C, PLAU and COL18A1 during senescence, while matched proteomic measurements were analysed to evaluate corresponding changes in secreted protein abundance. Additional transcriptomic datasets representing diverse stress conditions, including ionising radiation, etoposide treatment, oncogene-induced senescence, oxidative stress, hypoxia, and endoplasmic reticulum stress, were analysed to evaluate the behaviour of the EN526– CDKN2C regulatory axis across senescence and non-senescence contexts (Fig. 4f–k; GEO: GSE179810, GSE160356, GSE103938, GSE104666, GSE139963, GSE125218 ^31–36)^. Detailed information on all publicly available samples analysed in this study, including accession numbers, and experimental conditions, is provided in Supplementary Table S1.

**Figure 1.**
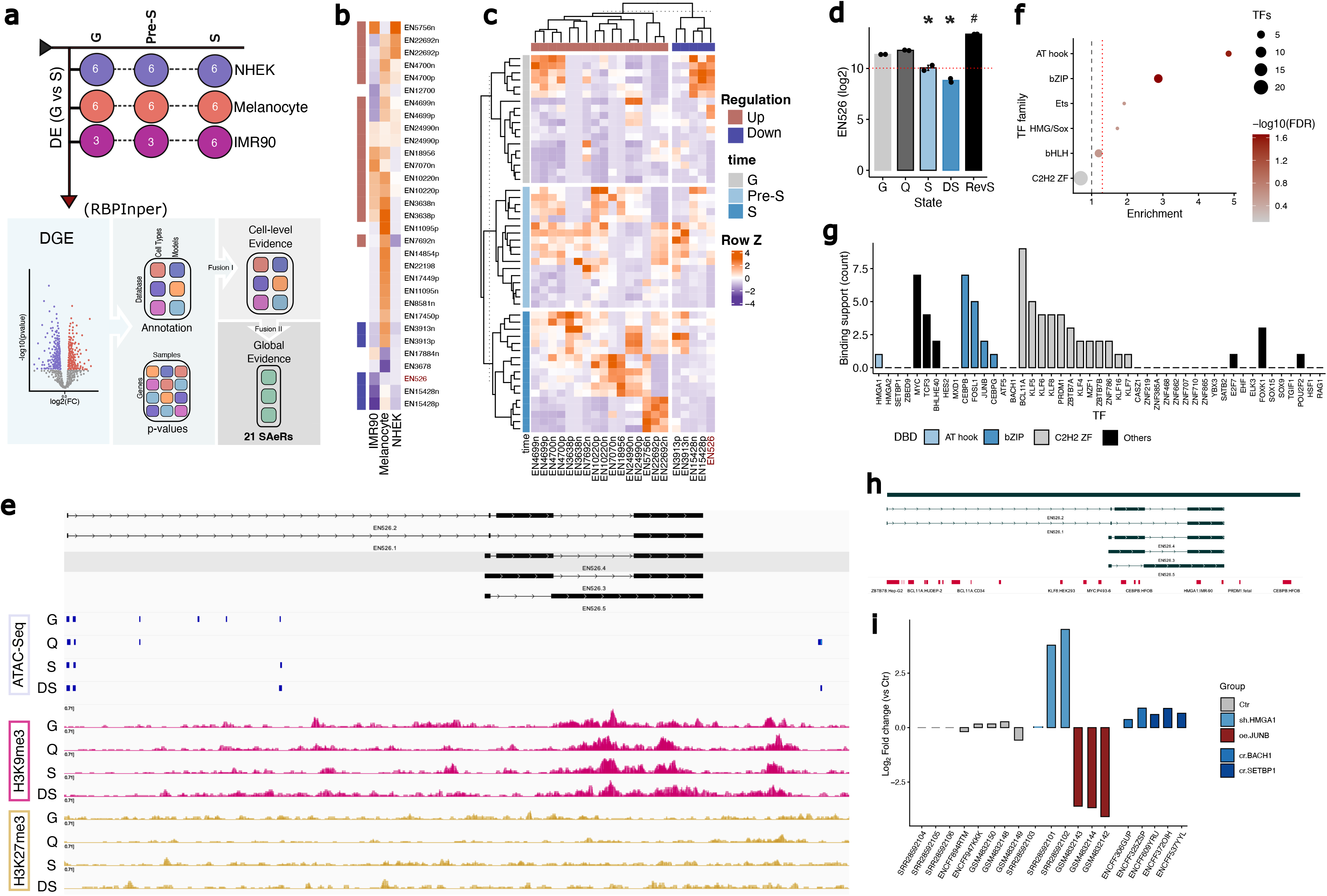
Identification and regulatory context of senescence-associated enhancer RNAs. **(a)** Schematic overview of the analytical framework used to identify senescence-associated enhancer RNAs (SAeRs). Differential expression analysis was performed across irradiation-induced senescence datasets from primary human fibroblasts (IMR90), keratinocytes (NHEK), and melanocytes using limma, followed by cross-cell-type integration with RBPInper to derive a consensus set of SAeRs. Numbers within circles indicate biological samples analysed for each cell state and cell type. **(b)** Heatmap showing log2 fold change of differential expression of identified SAeRs across IMR90, melanocytes, and NHEK cells, comparing growing (G) and senescent (S) states (Data: E-MTAB-5403 ^6^). **(c)** SAeR expression across time-resolved states (growing (G), pre-senescent (Pre-S), senescent (S)), in the same dataset analysed in (a–b). Samples are grouped by state and hierarchically clustered within each state to reveal similarities and heterogeneity in eRNA expression while preserving temporal ordering. Within **(b)** and **(c)**, red text indicates the prioritised SAeR. **(d)** Expression of EN526 across cellular states in BJ fibroblasts, including growing (G, n=2), quiescent (Q, n=2), senescent (S, n=2), deep senescent (DS, n=2), and senescence reversal (RevS, n=2) (Data: GSE133292 ^25^). Statistical significance was determined using limma empirical-Bayes analysis with FDR correction; *FDR < 0.05 vs G; #FDR < 0.05 vs DS. **(e)** Genomic view of the EN526 locus showing transcript models alongside ATAC-Seq signal and histone modification profiles (H3K9me3 and H3K27me3) across cellular states (growing (G); senescent (S); quiescent (Q); deep senescent (DS)); n=2 biological replicates per assay per state (Data: GSE133292 ^25^). **(f)** Enrichment analysis of DNA-binding domain families among transcription factors negatively correlated with EN526 expression in E-MTAB-5403 ^6^ across all samples (r ≤ −0.5). DBD-family enrichment was assessed using Fisher’s exact test with Benjamini– Hochberg correction for multiple testing. Dot size represents the number of transcription factors per family, and colour indicates the strength of −log10(FDR). **(g)** Distribution of ReMap-called transcription factor ChIP-Seq peaks across the EN526 locus ^26^, grouped by DNA-binding domain class. (**h**) EN526 locus annotated with selected ReMap-called transcription factor ChIP-Seq peaks from (g). **(i)** Relative EN526 expression following CRISPRi-mediated depletion of BACH1 (cr.BACH1) and SETBP1 (cr.SETBP1) in K526 from ENCODE RNA-Seq datasets, shRNA-mediated depletion of HMGA1 (sh.HMGA1) in KYSE-30 cells (GEO: GSE263439) ^28^, and JUNB overexpression (oe.JUNB) in A375 cells (GEO: GSE159539) ^29^. Bars represent expression log fold change relative to matched control cells; each bar represents a single sample.

**Figure 2.**
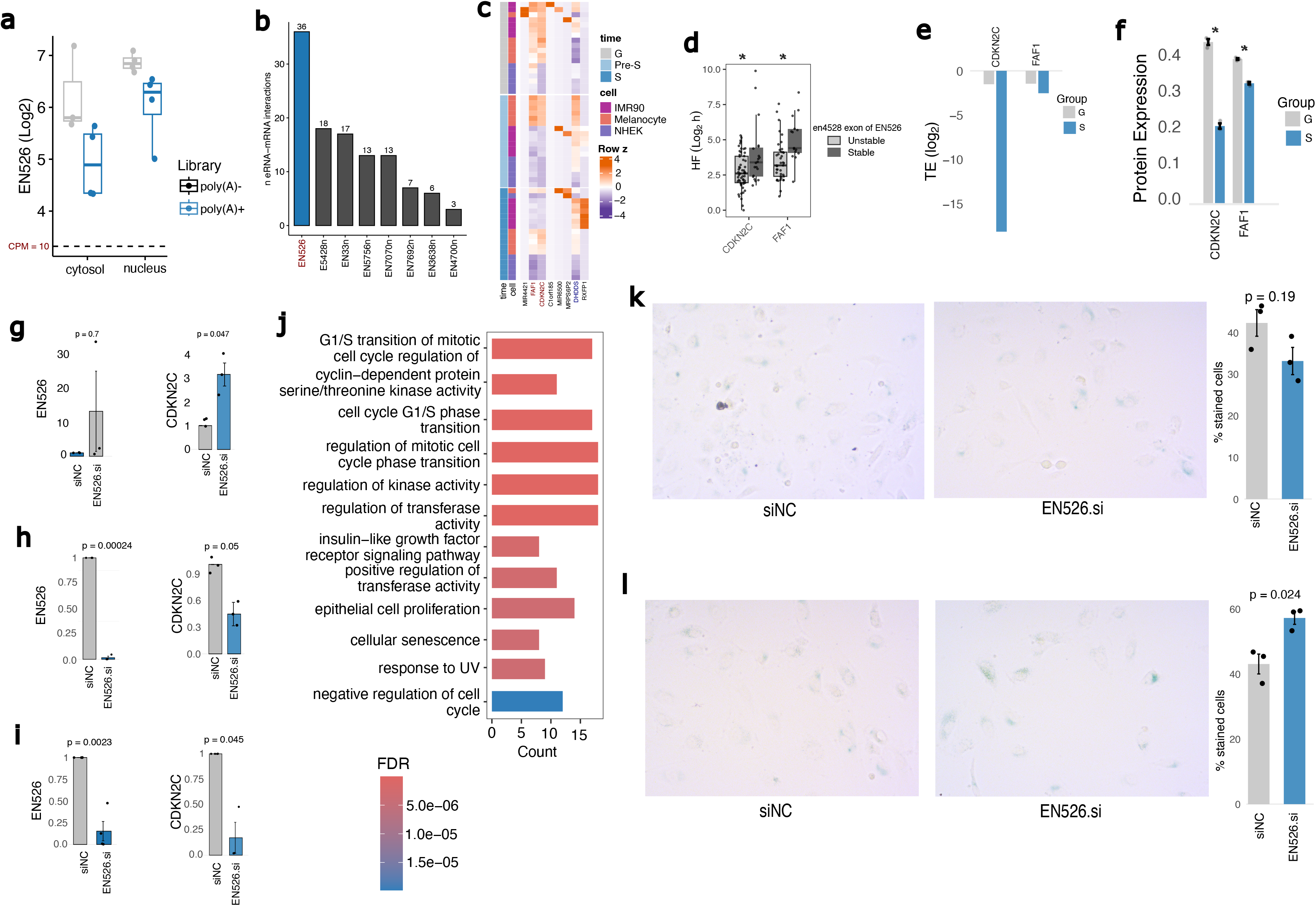
Post-transcriptional regulatory activity of EN526 and functional effects on senescence phenotypes. **(a)** EN526 expression in cytoplasmic and nuclear fractions across poly(A)-positive, poly(A)-negative, and total RNA-Seq libraries from eRNAkit resource (n=4 cell lines). **(b)** Number of high confidence (>2 samples) eRNA–mRNA interactions identified for SAeRs in the eRNAkit resource, with EN526 highlighted. **(c)** Heatmap showing expression of EN526 target mRNAs across IMR90, melanocytes, and NHEK cells in growing (G, n=15), pre-senescent (Pre-S, n=15), and senescent (S, n=18) states (Data: E-MTAB-5403 ^6^). **(d)** Distribution of mRNA half-lives (HF, log2 hours) for EN526 target transcripts (CDKN2C and FAF1), stratified by EN526 stability class in the Actinomycin D time-course datasets from the eRNAkit resource. * Indicates a significant difference in modelled mRNA half-life between the stable and unstable groups (FDR < 0.05), as determined in our previously published analysis ^24^. **(e)** Translation efficiency (TE, log2) of CDKN2C and FAF1 in growing (G; RNA-Seq n=2, Ribo-Seq n=3) and senescent (S; RNA-Seq n=3, Ribo-Seq n=3) states from GSE227766 ^30^. TE was calculated from aggregated samples as the log₂ ratio of Ribo-Seq to RNA-Seq reads for each gene. **(f)** Protein expression levels of CDKN2C and FAF1 in growing (G, n=3) and senescent (S, n=3) states from the same dataset as **(e)**. * Indicates p<0.05, two-sided t-test. Protein expression is shown as normalised TMT reporter-ion intensity (arbitrary units). **(g-i)** Relative expression levels of EN526 and CDKN2C measured by RT-qPCR following **(g)** siRNA-mediated knockdown of EN526 (EN526.si) compared to non-targeting control (siNC) for 24 hours, **(h)** 48 hours post-transfection or (**i**) pretreatment with actinomycin D for 30 minutes before siRNA-mediated knockdown of EN526 (EN526.si) compared to non-targeting control (siNC) at 2 hours. GAPDH mRNA was used as reference gene. Data are shown as mean ± SEM from at least three independent experiments. Indicated p-values were calculated using two-sided t-tests, except in (**i**), where a one-sided t-test was used to test the prespecified directional hypothesis that EN526 depletion reduces CDKN2C expression following actinomycin D pre-treatment. (**j**) Gene ontology enrichment analysis of CDKN2C protein interaction network (BioGRID database), showing top 12 enrichment for biological process terms with reduced redundancy via semantic similarity filtering. (**k-l**) The expression of senescence-associated ß-galactosidase (SA-ß-gal) after siRNA-mediated knockdown of EN526 (EN526.si) compared to non-targeting control (siNC) for hours 24 hours (**j**) or (**k**) 12 hours pre-siRNA transfection followed by 24 hours treatment with doxorubicin to induce senescence. Left panel shows representative images for EN526.si and non-targeting control cells. Right panel quantifies percentage of stained cells (mean ± SEM from three independent experiments). Indicated p-values were calculated using two-sided t-tests.

Raw reads corresponding to these datasets were obtained from the Sequence Read Archive (SRA) and processed using our established pipeline ^37^. Briefly, reads were quality filtered (Q < 20) and adapter trimmed using Trimmomatic (v0.39) ^38^, then aligned to the human reference genome (hg38) using HISAT2 (v2.1.0) with default parameters ^39^. Gene level and eRNA counts were quantified using HTSeq (v0.11.1)^40^ against our transcript-resolved eRNA GTF ^41^ or human GRCh38 annotation (Ensembl v111 release). Expression levels were normalised using counts per million (CPM), with gene and eRNA expression quantified and normalised independently to account for differences in feature abundance.

Translation efficiency (TE) was calculated as the log₂ ratio of ribosome profiling (Ribo-Seq) reads to RNA-Seq reads for each gene. Protein abundance values were obtained directly from the processed quantitative proteomics tables provided by the original study in the associated supplementary materials. ChIP–Seq datasets were analysed using pre-processed bigwig signal tracks downloaded from GEO accessions above. For ATAC–Seq datasets, precomputed peak calls provided by the original studies were used for downstream analyses.

### 2.2 Integration of transcriptomic datasets to define recurrent senescence-associated features

Differential expression (DE) analysis using limma was performed separately for each cell type for eRNAs and mRNAs, comparing proliferating cells with senescent or pre-senescent conditions ^42^. For each comparison, p-values were integrated across cell types using RBPInper (default parameters) ^37^, with eRNAs, mRNAs, and transcription factors analysed separately. Features showing significant global integration and consistent fold-change direction in at least two of three cell types were defined as recurrent.

### 2.3 Identification of candidate transcriptional repressors of EN526

TFs identified as recurrently dysregulated through the integrated analysis described above were examined as potential repressors of EN526. We focused on TFs showing recurrent upregulation across senescence models and correlated their expression with EN526 across samples using Pearson correlation. Across 1,610 TFs tested, 196 showed strong negative correlation (r ≤ −0.5), 294 showed strong positive correlation (r ≥ 0.5), and 1,120 showed weaker correlations (−0.5 < r < 0.5). TFs showing strong negative correlation were retained as candidate repressors. We then assessed enrichment of DNA-binding domains among these candidates to identify overrepresented TF families. DNA-binding domain (DBD) annotations were obtained from curated transcription factor classification resources ^43,44^. For each DBD family, a 2×2 contingency table comparing the number of candidate TFs carrying that domain with the remaining TFs in the background set was constructed, and enrichment was assessed using Fisher’s exact tests. Resulting p-values were adjusted for multiple testing using the Benjamini–Hochberg method. Finally, we assessed evidence for direct binding at the EN526 locus by interrogating ReMap ChIP-Seq datasets ^26^ for peaks overlapping the EN526 enhancer region. We also examined EN526 expression in publicly available transcription-factor perturbation RNA-Seq datasets, including CRISPRi-mediated depletion of SETBP1 and BACH1 in K562 cells ^27^, depletion of HMGA1 in KYSE-30 cells, and overexpression of JUNB in A375 cells ^28,29^.

### 2.4 eRNA annotation, interaction, and stability datasets

The eRNA annotation, catalogue of high-confidence eRNA–mRNA interactions, and eRNA stability effect data used in this study were derived from the eRNAkit resource, as previously described ^23,24^. Briefly, the eRNA–mRNA interaction catalogue was generated from the original datasets underlying the KARR-Seq ^45^, RIC-Seq ^46^ and PARIS ^47^ method papers (GEO accessions: GSE166155, GSE190214, GSE127188 and GSE74353). Raw reads were downloaded from NCBI GEO, quality-filtered and re-aligned using a standardised workflow, and chimeric alignments linking eRNAs with annotated mRNAs were parsed and aggregated to produce a curated, high-confidence and reproducible interaction catalogue. eRNA stability effect annotations were obtained from the precomputed kinetic modelling analysis reported in the associated study ^24^. Localisation profiles for EN526 were obtained from the eRNAkit datasets, which incorporate ENCODE fractionation RNA-Seq experiments. EN526 expression across nuclear and cytoplasmic fractions and across poly(A)+, poly(A)–, and total RNA libraries was extracted to examine its subcellular distribution across 13 human cell lines, as previously described ^23^.

In the eRNAkit resource, en4528 and en4529 represent two adjacent enhancer genomic coordinates, whereas subsequent transcript reconstruction demonstrated that these DNA regions overlap exonic regions of the same EN526 transcriptional unit, which generates five processed isoforms ^41^. Accordingly, eRNA–mRNA interactions were recomputed from the raw chimeric-junction data using the reconstructed EN526 annotation (Fig. 2b). For the previously established coordinate-based kinetic analysis shown in Fig. 2d, en4528 is retained and explicitly identified as an exonic region of EN526.

### 2.5 Interaction network interrogation and functional enrichment analysis

Protein–protein interaction data for CDKN2C were obtained from the BioGRID database ^48^. Experimentally validated interactions were extracted and filtered to retain interactions supported by more than two independent sources, generating an extended CDKN2C interaction network. Gene ontology enrichment analysis was then performed on the resulting network. We employed the ClusterProfiler R package ^49^ along with the human Bioconductor annotation database (org.Hs.eg.db) to identify enriched biological processes at FDR < 0.05, reducing redundancy via semantic similarity analysis ^50^.

Functional interaction networks for CDKN2C, PLAU, and COL18A1 were interrogated using the STRING database through the web interface ^51^. The analysis was used to evaluate potential functional associations between these proteins across curated pathways, protein– protein interactions, and other evidence channels. Genes identified within the resulting STRING interaction network were extracted and carried forward for downstream enrichment analysis. Functional enrichment of the STRING-derived gene set was performed using Enrichr ^52^ via the online platform to identify overrepresented GO biological processes.

### 2.6 Cell culture

Primary human umbilical vein endothelial cells (HUVEC) were obtained from PromoCell and certified mycoplasma-free. HUVEC cells were maintained in Endothelial Cell Growth Medium (PromoCell) supplemented with Endothelial Cell Growth Medium SupplementPack (PromoCell). Cells were cultured at 37°C in a humidified incubator with 5% CO₂. siRNA inhibitors were transfected using Lipofectamine RNAiMAX (Thermo Fisher Scientific, UK), as previously described ^24^. Details of siRNA sequences and primers used in this study are listed in Table S2.

### 2.7 RNA extraction and RT-qPCR

Total RNA was extracted from EN526.si, or siNC transfected cells using TRIzol reagent (Thermo Fisher, UK) according to the manufacturer’s instructions. cDNA was synthesised from 500⍰ng of total RNA using the iScriptTM cDNA Synthesis Kit (Bio-Rad, UK). Quantitative PCR was performed on a CFX96 Real-Time PCR Detection System (Bio-Rad, UK). The ΔΔCT analysis method was used, normalised to the GAPDH mRNA as reference gene. The Primer sequences and genomic coordinates are provided in Table S2.

### 2.8 SA-β-galactosidase assay

Cells were fixed, washed three times in PBS, and stained overnight at 37⍰°C using the Senescence beta-Galactosidase Staining Kit (Cell Signalling #9860), according to the manufacturer’s instructions. Cells were seeded at 3.5 × 10^4^ (24 well plates) or 1.15 x 10^5^ (6 well plates) cells per well prior to transfection and treatment. For assessment of basal effects of EN526 depletion, cells were transfected with EN526.si or siNC control siRNA and stained after 24 h. To evaluate the effect of EN526 depletion on stress-induced senescence, cells were first transfected with siRNA for 12 hours and then treated with doxorubicin (250 nM) for an additional 24 hours prior to staining. SA-β-galactosidase-positive cells were manually counted relative to the total number of cells per field using the multipoint tool in ImageJ ^53^. We quantified senescence by analysing 13 image fields per biological replicate and calculating the percentage of stained cells on a per-image basis. Image-level measurements were then averaged within each biological replicate prior to statistical testing.

### 2.9 DAPI-based nuclear morphology analysis

For nuclear morphology analysis, cells were fixed and stained with DAPI following either 24 hours of EN526 knockdown or pre-treatment with EN526 siRNA (12 hours) followed by doxorubicin exposure (24 hours), as described above. Images were acquired using an EVOS FL Auto 2 Imaging System (10× objective), and nuclei were segmented using a custom Python-based image analysis pipeline https://github.com/AneneLab/Eden “morphology”. Nuclear area and circularity were quantified for each nucleus, and per-image summaries were calculated prior to aggregation at the biological replicate level for statistical analysis.

### 2.10 Scratch wound assay

For wound healing assays, a uniform scratch was introduced into confluent monolayers of cultured cells using a 200μl pipette tip, followed by incubation in Optimem reduced-serum media. Images were captured at defined intervals over 48⍰h using an EVOS FL Auto 2 Imaging System (10× objective). The number of cells remaining attached within the imaged field was manually counted using the multipoint tool in ImageJ.

### 2.11 MTT cell viability assay

Cells were seeded in 96-well plates at 2 × 10⁴ cells per well. MTT reagent (0.5 mg/ml) was added and incubated for 3–4 h at 37 °C. Formazan crystals were dissolved in DMSO, and absorbance was measured at 540 nm with background correction at 690 nm. Values were normalised to the 690 nm signal and expressed relative to time 0.

### 2.12 Cytokine array assay

The Proteome Profile Human Angiogenesis Array kit, (R&D Systems) was used according to the manufacturer’s instruction. Briefly, conditioned media from serum and growth factor starved HUVEC (2⍰×⍰10^5⍰^ cells/ml in Optimem reduced-serum media) transfected with EN526.si or siNC siRNAs at 37⍰°C for 48⍰hours was collected and centrifuged to remove cellular debris at 500g. After blocking of non-specific binding at room temperature antibody array membranes were incubated with conditioned media at 4⍰°C overnight then incubated with diluted horseradish peroxidase-conjugated streptavidin (room temperature, 30⍰min) and visualised by ChemiDoc Imaging System (BioRad) Quantification was carried out using ImageJ.

### 2.13 Statistical analysis

Unless otherwise stated, graphical data represent the mean ± standard error of the mean (SEM) from at least three independent experiments. Differences between groups were assessed using Student’s t-test, Wilcoxon rank-sum test, or one-way ANOVA, as specified in the figure legends. Statistical significance was defined as p⍰<⍰0.05 for single comparisons and as FDR⍰<⍰0.05 for multiple testing.

## 3. Results

### 3.1 Recurrent enhancer RNA dysregulation defines the senescent state across cell types

To determine whether eRNAs are recurrently altered during senescence, we analysed a publicly available time-resolved RNA-Seq dataset in which senescence was induced by irradiation across three primary human cell types: fibroblasts, keratinocytes (NHEK), and melanocytes (Fig. 1a, Table S1). This design enabled identification of eRNAs associated with the fully established senescent state while also allowing their behaviour to be examined during the transition into senescence. Given our previous evidence that polyadenylated eRNAs are enriched among transcripts that engage in post-transcriptional regulatory interactions and show defined subcellular localisation ^24^, we prioritised poly(A)-selected RNA-Seq libraries. Importantly, poly(A)-based sequencing still captures eRNA transcription broadly, allowing detection without strong bias while favouring transcripts more likely to have functional regulatory roles. We first analysed 2,566 annotated eRNA loci ^41^ that were sufficiently expressed across the samples for differential expression analysis comparing proliferating controls with senescent samples within each cell type. At FDR < 0.05, this identified 42 differentially expressed eRNAs in fibroblasts (20 upregulated, 22 downregulated), 65 in NHEK (57 upregulated, 8 downregulated), and 56 in melanocytes (47 upregulated, 9 downregulated). Differential expression results were then integrated across cell types using RBPInper, which integrates the underlying p-values, to identify recurrent signals independent of cellular background ^37^. This analysis identified 21 eRNAs consistently dysregulated at the fully senescent stage (16 upregulated and 5 downregulated, Fig. 1b-c), hereafter referred to as senescence-associated eRNAs (SAeRs). Across the SAeR set, strand-resolved eRNA transcripts originating from the same enhancer locus generally changed in the same direction (12 upregulated vs 4 downregulated, Fig. 1c), indicating coordinated enhancer reprogramming during senescence. Three upregulated SAeRs were represented by a single strand-resolved transcript (EN5756n, EN7070n, and EN7692n), while two SAeRs (EN18956 and EN526) correspond to transcripts whose strand orientation is defined by the reference annotation ^41^.

Having defined this integrated SAeR set, we next examined their expression across intermediate time points to determine how they behave during senescence progression. Most SAeRs exhibited stable ON/OFF-like dynamics, with transcripts that were induced in the final senescent state already elevated at earlier stages, and transcripts that were repressed remaining low throughout the trajectory (Fig. 1c). A smaller subset showed gradual or transient changes, suggesting that these may reflect regulatory adjustments occurring during the establishment rather than the maintenance of senescence (Fig. 1c). Together, these analyses indicate that a core group of eRNAs is reproducibly altered at the fully senescent state, consistent with coordinated enhancer reprogramming during senescence.

### 3.2 EN526 repression during senescence involves stress-responsive transcription factors

To determine whether SAeRs contribute to senescence beyond reflecting enhancer transcriptional activity, we next focused on EN526 ( chr1:50977923-51021398, hg38), one of the five downregulated SAeRs identified across the three cell types. While enhancer reprogramming is a well-established feature of senescence, whether eRNAs themselves influence the phenotype through altered post-transcriptional regulatory mechanisms remains unclear. To examine this possibility, we first assessed structural features and existing annotations across the 21 SAeRs. This analysis identified EN526 as a locus with several properties suggesting potential RNA-level regulatory activity. First, EN526 overlaps an eRNA we previously reported to influence target mRNA stability and translation efficiency ^24^. Second, unlike most SAeRs, the EN526 locus is processed and generates multiple isoforms. Specifically, it encodes five spliced transcripts ranging from 4.15 kb to 8.90 kb in length (EN526.1: 4,154 bp; EN526.2: 7,351 bp; EN526.3: 7,816 bp; EN526.4: 7,497 bp; EN526.5: 8,895 bp) ^41^. Based on these features, we selected EN526 for further investigation as a candidate eRNA with potential post-transcriptional function altered in senescence.

To confirm that EN526 dysregulation is reproducible across senescence models, we examined its expression in an independent RNA-Seq cohort generated in BJ fibroblasts spanning multiple cellular states, including proliferating, quiescent, senescent, deep senescent, and senescence reversal, in which senescence was induced by bleomycin ^25^. EN526 expression was reduced in both senescent states relative to proliferating and quiescent cells (Fig. 1d), indicating that its repression is associated with senescence rather than cell-cycle arrest alone. EN526 expression increased upon senescence reversal, consistent with repression that tracks the senescent state. To examine the chromatin context of the EN526 locus, we analysed ATAC-Seq and histone modification profiles across the same BJ cell states. ATAC-Seq signal was detected at discrete sites within the locus, and when present these peaks occurred at similar positions across all states (Fig. 1e), indicating stable local accessibility rather than widespread chromatin opening in this model. Consistent with this, the repressive histone marks H3K9me3 and H3K27me3 showed largely similar distributions across the locus in all conditions (Fig. 1e). Together, these profiles suggest that the chromatin environment of EN526 remains structurally stable during senescence in BJ cells. Notably, the presence of these repressive marks does not appear to preclude local accessibility or eRNA transcription at the locus, suggesting that enhancer activity at EN526 can persist within chromatin domains carrying broadly repressive signatures when accessibility is maintained at specific sites.

Given the absence of major differences in repressive chromatin features across cell states, we next examined whether transcription factor (TF) binding at the EN526 locus could partly explain the observed changes in EN526 expression. We repeated the differential expression analysis followed by RBPInper integration, this time focusing on mRNAs. We then selected TFs that were consistently upregulated across the senescence samples, reasoning that increased abundance of repressive TFs could contribute to repression through direct binding at the locus. Correlating the expression of these TFs with EN526 across samples (r ≤ −0.5) identified 47 candidate regulators. To determine whether these TFs share structural features associated with potential regulatory activity at this locus, we examined enrichment of their DNA-binding domains (DBDs). This analysis revealed significant enrichment of AT-hook (n = 3) and bZIP (n = 5) transcription factor families (FDR < 0.05; Fig. 1f). Both families are well known mediators of stress-responsive transcriptional programs and often act as repressive factors. To assess whether these candidates show direct evidence of binding at the EN526 locus, we interrogated the ReMap ChIP–Seq database ^26^. Several of the predicted TFs displayed binding support on the EN526 region across multiple cell types (Fig. 1g), with AT-hook and bZIP factors showing the most frequent binding signals distributed along the locus (Fig. 1h), consistent with a potential role for these TF families in the regulation of EN526 during senescence.

To assess whether putative TFs identified by our analysis could influence EN526 expression, we examined publicly available TF-perturbation datasets for representative AT-hook and bZIP factors. We initially selected SETBP1 (AT hook) and BACH1 (bZIP) because, despite being identified by the correlation and DBD-enrichment analyses, neither had available ChIP–Seq evidence of binding at the EN526 locus (Fig. 1g). This provided an orthogonal test of candidates for which the absence of binding evidence could reflect limited experimental coverage rather than absence of regulatory activity. Depletion of both TFs was associated with increased EN526 expression relative to control conditions in K562 cells (Fig. 1i) ^27^, providing additional support for the candidate-TF analysis. We then examined independent perturbations of HMGA1 (AT hook) and JUNB (bZIP), two candidate TFs with binding evidence at the EN526 locus. EN526 was undetectable in all KYSE-30 control samples but became detectable in two of three HMGA1-depleted samples, whereas JUNB overexpression in A375 cells strongly repressed EN526, reducing its expression by an average of 13.7-fold relative to controls (Fig. 1i) ^28,29^. Together, the senescence-associated upregulation and negative correlation of candidate TFs with EN526, their binding profiles across the locus, and the directionally consistent effects observed across independent depletion and overexpression experiments support a model in which stress-responsive TFs could contribute to EN526 repression, without requiring major changes in the surrounding repressive epigenetic landscape. These findings provide upstream regulatory context consistent with the observed repression of EN526 during senescence and suggest that the additive effect of multiple candidate TFs upregulated during senescence may reduce activity of the EN526 enhancer. Additional contributions from higher-order chromatin organisation, known to be remodelled during senescence ^7–9^, could further reinforce repression at the locus.

### 3.3 EN526 engages eRNA–mRNA interactions and post-transcriptionally regulates CDKN2C

Having defined the transcriptional repression of EN526 during senescence, we next shifted to examining whether the loss of EN526 RNA itself contributes to the senescent phenotype through altered post-transcriptional regulatory mechanisms. A key predictor of such a model is that the eRNA should normally be detectable outside the nucleus and engage in direct eRNA-mRNA interactions with target transcripts. Indeed, analysis of ENCODE localisation datasets showed that EN526 is detectable in cytoplasmic fractions from both K562 and HepG2 cells, with no strict dependence on poly(A) status for localisation (Fig. 2a). Consistent with this observation, interrogation of our curated eRNA-mRNA interaction datasets ^23^ revealed that EN526 exhibits approximately twofold more physical eRNA–mRNA interactions than the next most highly connected SAeRs (36 unique mRNAs, Fig. 2b). Using the same differential expression and RBPInper integration framework described above, we next examined the expression of EN526 target mRNAs across the senescence datasets. Two targets, CDKN2C and FAF1, were significantly reduced in senescence, whereas DHDDS was upregulated (Fig. 2c). Given the consistent reduction of CDKN2C and FAF1 together with EN526 and our previous evidence that eRNAs can stabilise mRNA targets independent of transcription ^24^, we next focused on these targets.

Kinetic modelling of actinomycin D time-course datasets using expression measured over the region formerly designated en4528 ^24^, now resolved as an exonic region of EN526 ^41^, associated this region with increased stability of CDKN2C and FAF1 mRNAs (Fig. 2d). This relationship is consistent with a stabilising role for EN526 and predicts that its repression during senescence would reduce CDKN2C and FAF1 expression. Together, these findings suggest that the reduced CDKN2C and FAF1 levels observed in senescence may be associated with loss of EN526 stabilising effects. Although both genes lie in genomic proximity to the EN526 locus, we previously demonstrated that such eRNA regulation can occur independently of transcription, consistent with a post-transcriptional mechanism ^24^. Changes in mRNA stability are expected to influence translation efficiency and ultimately protein output. We therefore examined translation efficiency (TE), calculated as the log₂ ratio of Ribo-Seq to RNA-Seq reads, and matched proteomic abundance measured by mass spectrometry for CDKN2C and FAF1 in senescent cells compared with proliferating controls. Consistent with this model CDKN2C showed a marked reduction in translation efficiency in senescent cells (Fig. 2e), accompanied by significantly reduced protein levels (p < 0.05; Fig. 2f). FAF1 exhibited a similar but weaker pattern, with slightly lower translation efficiency and significantly reduced protein levels (Fig. 2e-f).

To directly test whether EN526 regulates its target transcripts in primary cell model, we next perturbed EN526 using siRNA in primary endothelial cells, a well-established cell for studying stress-induced senescence phenotypes relevant to ageing and vascular biology. As siRNA activity is mediated through the RNA-induced silencing complex (RISC), this approach preferentially targets the cytoplasmic pool of the eRNA, enabling assessment of its potential post-transcriptional regulatory activity ^24^. Based on the stronger translation efficiency changes observed for CDKN2C, we focused subsequent analyses on this target. At 24 hours after siRNA treatment, EN526-directed siRNA induced a strong increase in CDKN2C expression (Fig. 2g), while EN526 levels themselves were not yet consistently reduced. By 48 hours, EN526 transcripts were effectively depleted, and the initial increase in CDKN2C expression was completely reversed (p < 0.05; Fig. 2h). Importantly, this regulatory effect was retained after actinomycin D pre-treatment (p < 0.05; Figure 2i), which blocks nascent RNA synthesis, excluding transcriptional feedback or secondary effects resulting from altered transcriptional programmes. Together, these observations demonstrate that perturbation of EN526 alters CDKN2C expression in a manner consistent with eRNA-mediated post-transcriptional regulation. As CDKN2C is a stress-responsive cell-cycle regulator implicated in senescence programs^54^, this EN526–CDKN2C axis provides a candidate mechanism through which eRNAs may influence stress-associated cellular phenotypes at the post-transcriptional level.

We next asked whether the EN526–CDKN2C post-transcriptional regulatory axis could reinforce the senescence phenotype through broader downstream networks. A simple prediction of this model is that protein interactors downstream of this eRNA–mRNA target would be enriched for pathways involved in cell-cycle regulation and stress-associated processes linked to senescence. To identify these downstream components, we extended the network by incorporating experimentally derived CDKN2C protein–protein interactions from the BioGRID database ^48^, retaining top 100 recurrent interactions supported by more than two independent sources. Subsequent gene ontology analysis of this extended network revealed significant enrichment for senescence-associated regulatory processes, including negative regulation of the cell cycle, regulation of cyclin-dependent kinase activity, G1/S phase transition, epithelial cell proliferation, cellular senescence, and cellular responses to stress such as UV signalling (Fig. 2j). These findings suggest that loss of EN526 during senescence, coupled with downstream destabilisation of CDKN2C, may reinforce the senescent state through broader regulatory networks governing cell-cycle control and stress-responsive signalling.

### 3.4 Loss of EN526-mediated post-transcriptional regulation phenocopies key senescence features

To determine whether EN526 depletion is sufficient to induce the senescence state, we first examined senescence-associated β-galactosidase staining following 24 hours of EN526 knockdown. Under these conditions, we observed no significant difference between EN526 knockdown and non-targeting control cells (Fig. 2k), indicating that EN526 loss alone is not sufficient to induce senescence. This is consistent with our earlier transcriptional analyses, which suggest that EN526 functions downstream of the primary stress-responsive senescence programme. We next asked whether prior depletion of EN526 alters cellular responses to senescence-inducing stress. We pre-treated cells with EN526 siRNA (12 hours) before exposing them to doxorubicin (plus 24 hours) to induce drug-associated senescence. Expectedly, we observed significantly higher senescence-associated β-galactosidase staining in EN526-depleted cells after 24 hours of doxorubicin treatment compared with control cells (p = 0.024; Fig. 2l). These observations indicate that EN526 loss amplifies senescent commitment under stress rather than triggering senescence autonomously.

Because senescence represents a stress-associated survival state in which cells remain viable despite permanent proliferative arrest, we next asked whether EN526 depletion alters cellular survival under stress conditions. To test this, we examined cellular behaviour under serum and growth factor starvation following EN526 knockdown. In scratch wound assays, we observed that both EN526-depleted and control cells underwent stress-induced cell loss, but a greater number of EN526-depleted cells remained attached at 48 hours (Fig. 3a-b). To quantify this effect independently, we repeated the experiment and measured cell viability using an MTT assay and confirmed that EN526-depleted cells have significantly greater viability at 48 hours than control cells (p = 0.0051; Fig. 3c). Together, these results indicate that EN526 depletion biases stressed cells toward a pro-survival state, consistent with EN526 loss sustaining the senescent phenotype.

**Figure 3.**
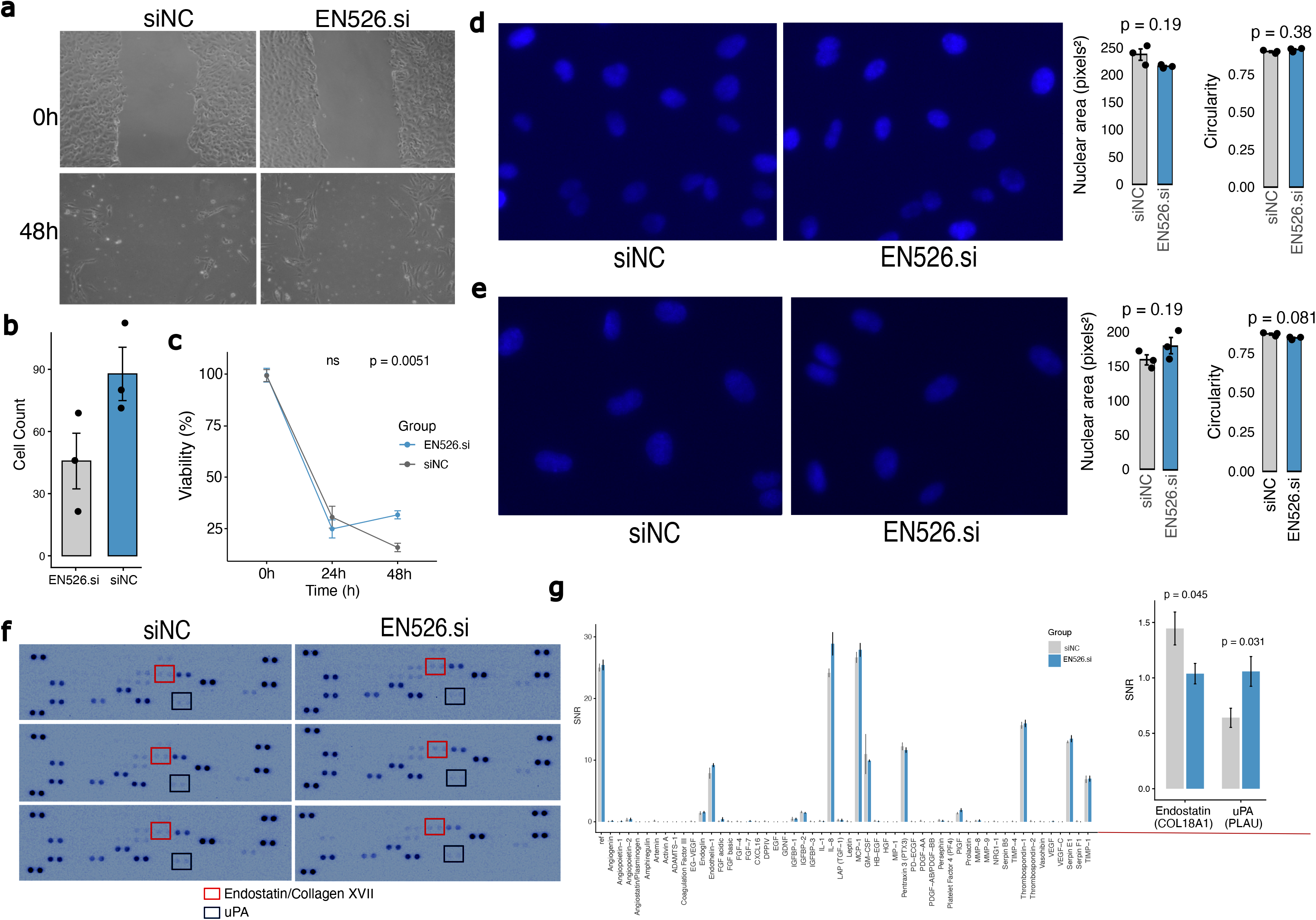
EN526 depletion modulates survival and secretory phenotypes without altering nuclear morphology. (**a-b**) Scratch wound assay assessing the effect of EN526 knockdown on cell survival compared to non-targeting control (siNC) 48 hours post-transfection, (**a**) representative images 0 and 48-hours post-scratch for EN526.si and siNC and (**b**) cell count over time (mean ± SEM from three independent experiments and 10 image views). (**c**) Cell viability measured by MTT assay at 0, 24, and 48 hours under no serum and growth factor supplement condition in EN526-depleted (EN526) and non-targeting control (siNC). (**d-e**) DAPI-based nuclei morphology analysis after siRNA-mediated knockdown of EN526 (EN526.si) compared to non-targeting control (siNC) for 24 hours (**d**) or (**e**) 12 hours pre-siRNA transfection followed by 24 hours treatment with doxorubicin to induce senescence. Left panel shows representative images for EN526.si and non-targeting control cells. Right panel quantifies nuclear area and nuclear circularity (mean ± SEM from three independent experiments). (**f-h**) Multiplex protein array analysis of secreted factors in conditioned media following siRNA-mediated knockdown of EN526 (EN526.si) compared to non-targeting control (siNC) for 48 hours. (**f**) All images of protein array membranes with Endostatin (COL18A1) and uPA (PLAU) dots highlighted. (**g**) Quantification of signal intensity across all assayed proteins; Endostatin (COL18A1) and uPA (PLAU) are enlarged for visualisation. Values shown in are signal-to-noise ratio (SNR) (mean ± SEM from three independent experiments). Across all panels, indicated p-values were calculated using two-sided t-tests, except in (g), where the Wilcoxon test was used. Across (a–f), indicated p-values were calculated using two-sided t-tests; in (g), p-values were calculated using the Wilcoxon rank-sum test.

**Figure 4.**
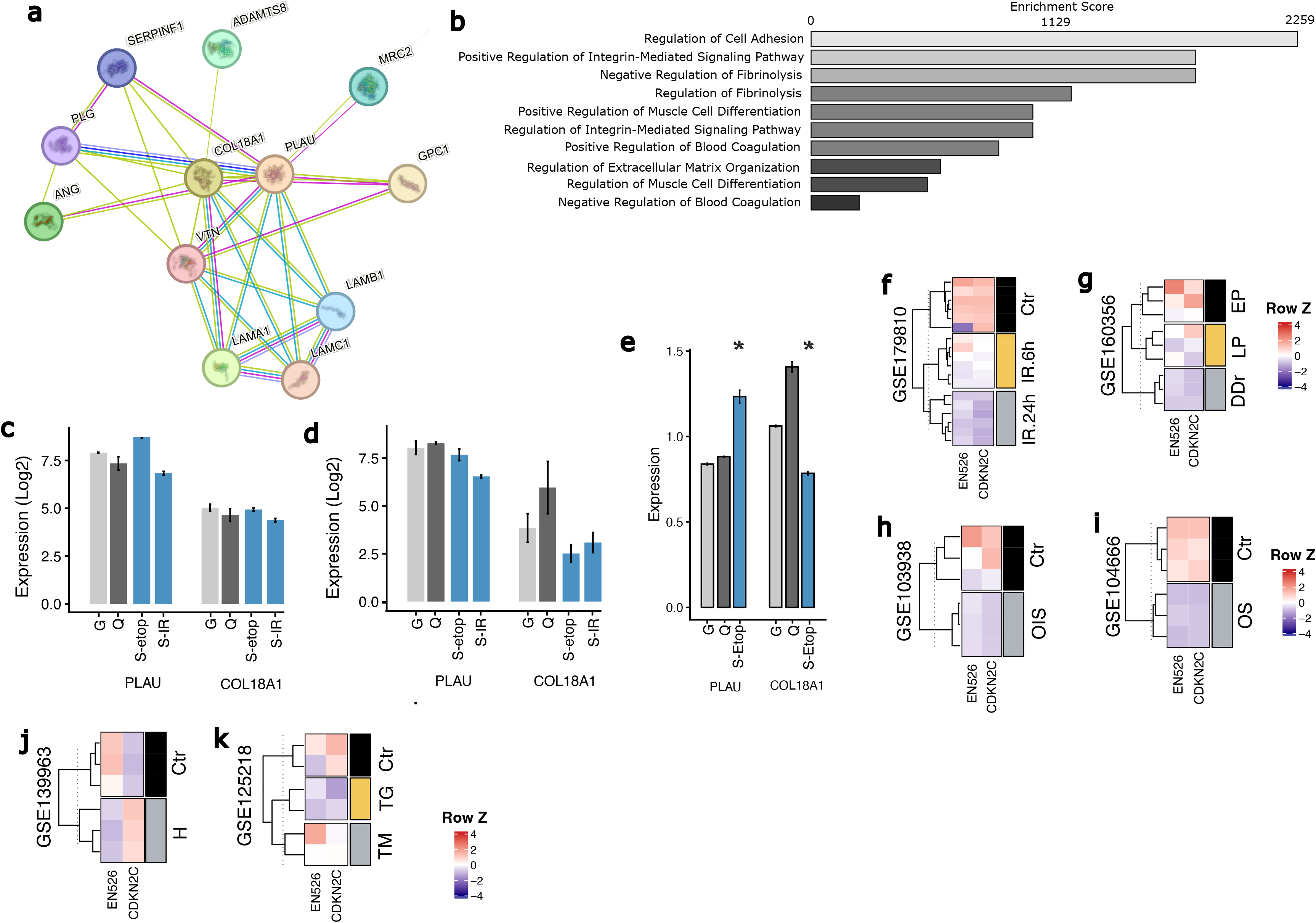
EN526-associated secretome changes and regulatory context across senescence and stress conditions. (**a-b**) Protein–protein and genetic interaction network analysis of PLAU and COL18A1 and associated interacting proteins derived from STRING analysis; (**a**) network plot (**b**) Gene ontology enrichment analysis of the extended interaction network showing Enrichr enrichment scores. (**c-e**) RNA expression (**c**), ribosome profiling (**d**), and protein expression (**e**) of COL18A1 (Endostatin) and PLAU (uPA) across cellular states including growing (G), quiescent (Q), etoposide-induced senescence (S-etop), and irradiation-induced senescence (S-IR) in GSE227766 ^30^; n=3 biological replicates per assay per group. In **(e)**, * indicates p<0.05, two-sided t-test. (**f–k**) Heatmaps showing expression of EN526 and CDKN2C across senescence and stress conditions; each heatmap observation represents an individual sample, with sample numbers and dataset details provided in Table S1 ^31–36^. (**f**) Expression of EN526 and CDKN2C following ionising radiation (IR) at 6 hours and 24 hours compared with control (Ctr) (Data: GSE179810; IR 6 h: EN526 p=0.00359, CDKN2C p=5.79×10⁻⁸; IR 24 h: EN526 p=2.32×10⁻⁶, CDKN2C p=2.60×10⁻⁶). (**g**) Expression of EN526 and CDKN2C across proliferating (EP), late passage (LP), and drug-induced senescence (DDr) states (Data: GSE160356; LP: EN526 p=0.335, CDKN2C p=0.294; DDr: EN526 p=0.0373, CDKN2C p=0.0129). (**h**) Expression of EN526 and CDKN2C in oncogene-induced senescence (OIS) compared with control (Ctr) (Data: GSE103938; EN526 p=0.399, CDKN2C p=0.0273). (**i**) Expression of EN526 and CDKN2C under oxidative stress conditions (OS) compared with control (Ctr) (Data: GSE104666; EN526 p=0.00230, CDKN2C p=0.000738). (**j**) Expression of EN526 and CDKN2C under hypoxia (H) compared with control (Ctr) (Data: GSE139963; EN526 p=0.0131, CDKN2C p=0.000229). (**k**) Expression of EN526 and CDKN2C under endoplasmic reticulum stress conditions, including tunicamycin (TM) and thapsigargin (TG), compared with control (Ctr) (Data: GSE125218; combined ER-stress comparison: EN526 p=0.804, CDKN2C p=0.0398). Indicated p-values were calculated using two-sided Welch’s t-tests on log₂-transformed expression values.

Senescence is frequently accompanied by large-scale transcriptional and chromatin reorganisation, which can manifest as changes in nuclear morphology, including increased nuclear size and irregular shape ^55^. We therefore asked whether EN526 depletion is associated with detectable alterations in nuclear architecture. To address this, we quantified nuclear area and circularity by DAPI staining under both basal and stress conditions. Following 24 hours of EN526 knockdown alone, we observed no significant differences in nuclear size or circularity compared with control cells (Fig. 3d). We next assessed whether pre-EN526 depletion modifies nuclear morphology in response to senescence-inducing stress. Cells were pre-treated with EN526 siRNA for 12 hours prior to doxorubicin exposure (24 hours), mirroring the conditions used to induce senescence. Under these conditions, we again observed no significant differences in nuclear area or circularity between EN526-depleted and control cells (Fig. 3e). Together, these results indicate that EN526 depletion does not induce detectable changes in nuclear morphology either at baseline or under stress, suggesting that the enhanced senescent phenotype is unlikely to be driven by gross alterations in nuclear architecture or large-scale genomic reorganisation.

Given the survival-associated senescent state, we next asked whether EN526 depletion alters the senescence-associated secretory phenotype (SASP). To test this, we profiled secreted proteins using a multiplex protein array comparing EN526-depleted and control cells. Among the 52 proteins assayed in the conditioned media, only two showed a significant difference (p < 0.05): uPA/PLAU was increased, whereas Endostatin/COL18A1 was reduced in EN526-depleted cells (Fig. 3f–g). PLAU promotes extracellular proteolysis through plasmin generation and activation of downstream proteases, while COL18A1 is a structural component of the basement membrane ^56,57^. Notably, IL-8, a canonical SASP factor, was also increased in EN526-depleted cells, although this change did not reach statistical significance. The observed increase in PLAU together with reduced COL18A1 suggests enhanced extracellular matrix remodelling, a common feature of the senescence-associated secretory phenotype. These changes are consistent with the altered survival and adhesion phenotypes observed above and support the role of EN526 loss in reinforcing senescence-associated cellular phenotypes.

To test whether the secretome changes observed following EN526 depletion might arise as a downstream consequence of CDKN2C regulation, we first examined potential physical and functional interactions between CDKN2C, PLAU, and COL18A1 using STRING network analysis ^51^. This analysis did not identify direct or high-confidence interactions linking CDKN2C to either PLAU or COL18A1 within the interaction network (Fig. 4a), suggesting that the observed secretome changes are unlikely to be mediated through CDKN2C-dependent pathways. These findings raise the possibility that EN526 may influence these proteins through mechanisms independent of its post-transcriptional regulation of CDKN2C. To place these proteins within a broader functional context, we next performed gene ontology enrichment analysis on the extended interaction network identified by STRING. We found significant enrichment for pathways associated with regulation of cell adhesion, integrin-mediated signalling, extracellular matrix organisation, and fibrinolysis (Fig. 4b). These processes are characteristic features of senescence-associated extracellular remodelling and cellular adhesion dynamics, suggesting that EN526-associated secretome changes occur within a broader network of proteins linked to senescence-related extracellular regulation.

We next asked at which regulatory level these proteins change during senescence (transcriptional, translational, or secretory level). Interrogation of the same senescence datasets used above for CDKN2C and FAF1 showed no significant differences in RNA abundance or ribosome occupancy for PLAU or COL18A1 across proliferating, quiescent, and senescent cells induced by etoposide or irradiation (Fig. 4c-d). These results indicate that these genes are not strongly regulated at the transcriptional or translational level during senescence. Examination of secretome proteomics data from the same dataset revealed a different pattern: secreted protein levels were significantly altered, with PLAU increased and COL18A1 decreased in senescent cells compared with proliferating or quiescent controls (Fig. 4e). These findings indicate that regulation of these factors during senescence occurs predominantly at the level of protein secretion or extracellular stability, rather than gene expression. Notably, the direction of these changes mirrors those observed following EN526 depletion, supporting a link between EN526 loss and remodelling of the senescence-associated secretome.

### 3.5 EN526-CDKN2C axis captures senescence states and predicts senescence commitment

A universal and reliable signature of cellular senescence remains elusive ^58^, with current strategies often relying on either single marker of limited specificity or complex multi-gene expression programs that are not grounded in mechanism and vary across senescence triggers ^5,59^. While upstream regulators such as TP53 and CDKN2A provide mechanistic insights, their interpretation as markers is often context dependent. Notably, senescence is almost universally characterised by extensive epigenetic reprogramming, including chromatin remodelling, histone modifications, and DNA methylation changes ^60–62^. However, the lack of a single, accessible assay that captures these diverse epigenetic alterations has limited their utility as universal biomarkers. To address the unmet need for robust and mechanistically grounded senescence markers, we propose the EN526-CDKN2C axis, which integrates transcriptional control, and post-transcriptional regulation. This two-component signature captures both the regulatory cause (stress responsive TFs) and effectors (CDKN2C) of the senescence program, with EN526 acting as a regulatory intermediary with enhancer-like properties and post-transcriptional influence.

To investigate the specificity and dynamic behaviour of the EN526-CDKN2C axis across stress contexts, we systematically profiled its expression across diverse datasets representing both confirmed and putative senescence-inducing conditions. First, we analysed gene expression following ionising radiation (IR), a potent genotoxic stressor known to eventually induce senescence. In this dataset, cells were harvested at 6 h and 24 h post-IR. Strikingly, as early as 6 hours after treatment, EN526 was robustly inhibited, accompanied by marked repression of CDKN2C, compared to controls (Fig. 4f). This rapid and coordinated shift in expression highlights the early engagement of this regulatory axis following stress exposure. Importantly, prior studies have demonstrated that IR triggers a DNA damage response cascade culminating in senescence over time, but this observation shows that the EN526-CDKN2C axis responds within hours, supporting its role as a sensitive early marker of senescence commitment.

To determine whether this signature is specific to bona fide senescence rather than general proliferative decline, we next analysed gene expression in cells subjected to etoposide-induced senescence, in parallel with early and late passage cells that were not senescent. The full axis was precisely coordinated in the etoposide condition, EN526 repression with concomitant repression of CDKN2C whereas no such pattern was observed in late passage cells (Fig. 4g). This finding is notable because late passage cells often exhibit some transcriptional drift or stress features, but are not terminally arrested, indicating that the axis specifically captures senescence rather than chronological passage alone. In oncogene-induced senescence (OIS) models driven by RAS overexpression, EN526 was reduced (-0.83 log fold change), although greater variability between samples in this dataset meant that this change did not reach statistical significance, whereas CDKN2C was repressed (Fig. 4h). This variability in EN526 may reflect the direct effects of oncogenic RAS signalling on enhancer activity.

We next turned to oxidative stress, induced by treatment with 200 µM H₂O₂ for 36 hours. Although senescence was not formally confirmed in this dataset, this dose and duration of H₂O₂ are widely reported to induce senescence phenotypes ^63^. Notably, we observed full coordination of the axis (Fig. 4i), strongly suggesting that it can detect cellular states that are en-route to senescence, even before classical hallmarks become evident. To test whether this axis is merely a general stress response signature, we next examined its behaviour under two non-senescent stress conditions. In hypoxia-induced stress, EN526 was again repressed, however, CDKN2C expression was induced, indicating partial activation of the axis (Fig. 4j). Similarly, in endoplasmic reticulum (ER) stress induced by unfolded protein response (UPR) activation, EN526 was not repressed, whereas CDKN2C was consistently repressed (Fig. 4k).

Together, these findings support a model in which the full coordination of the EN526-CDKN2C axis is a hallmark of bona fide or impending senescence. Partial activation of the axis in other stress conditions may reflect a modular regulatory mechanism, where EN526 serves as a key integrator linking stress-responsive transcriptional events to downstream post-transcriptional control, but only engages effector gene repression fully in the context of senescence. This layered regulation highlights the potential of the EN526-CDKN2C axis as a robust biomarker for senescence, with both diagnostic and prognostic applications in aging and stress related conditions.

### 3.6 Genetic associations link EN526-CDKN2C axis to aging-related traits

To explore the potential relevance of the EN526 enhancer locus in human aging and disease, we examined whether genetic variants within this regulatory region are associated with age-related traits. We intersected the genomic coordinates of the EN526 enhancer with single-nucleotide polymorphisms (SNPs) catalogued in genome-wide and phenome-wide association studies (GWAS/PheWAS) ^64^. This analysis identified multiple trait-associated SNPs overlapping the EN526 locus (Fig. 5a), with most of them in linkage disequilibrium with CDKN2C. The age-associated traits include red blood cell indices, glycated haemoglobin levels (HbA1c), sex hormone–binding globulin, renal cell carcinoma, and cardiovascular phenotypes such as JT interval, cataracts, pain (Fig. 5b). The presence of these variants within the EN526 regulatory region suggests that genetic variation at this enhancer may influence physiological pathways linked to aging and chronic disease through altered eRNA levels ^65^. Notably, the linkage of these variants to CDKN2C, a key effector within the EN526 regulatory axis identified here, further supports the potential functional relevance of this enhancer locus in age-associated biological processes. In addition to associations with age-related phenotypes and cancer risk, variants at the EN526 locus show a striking enrichment for associations with circulating protein levels (Fig. 5b), particularly secreted proteases, suggesting a broader role in systemic or post-transcriptional regulation.

**Figure 5.**
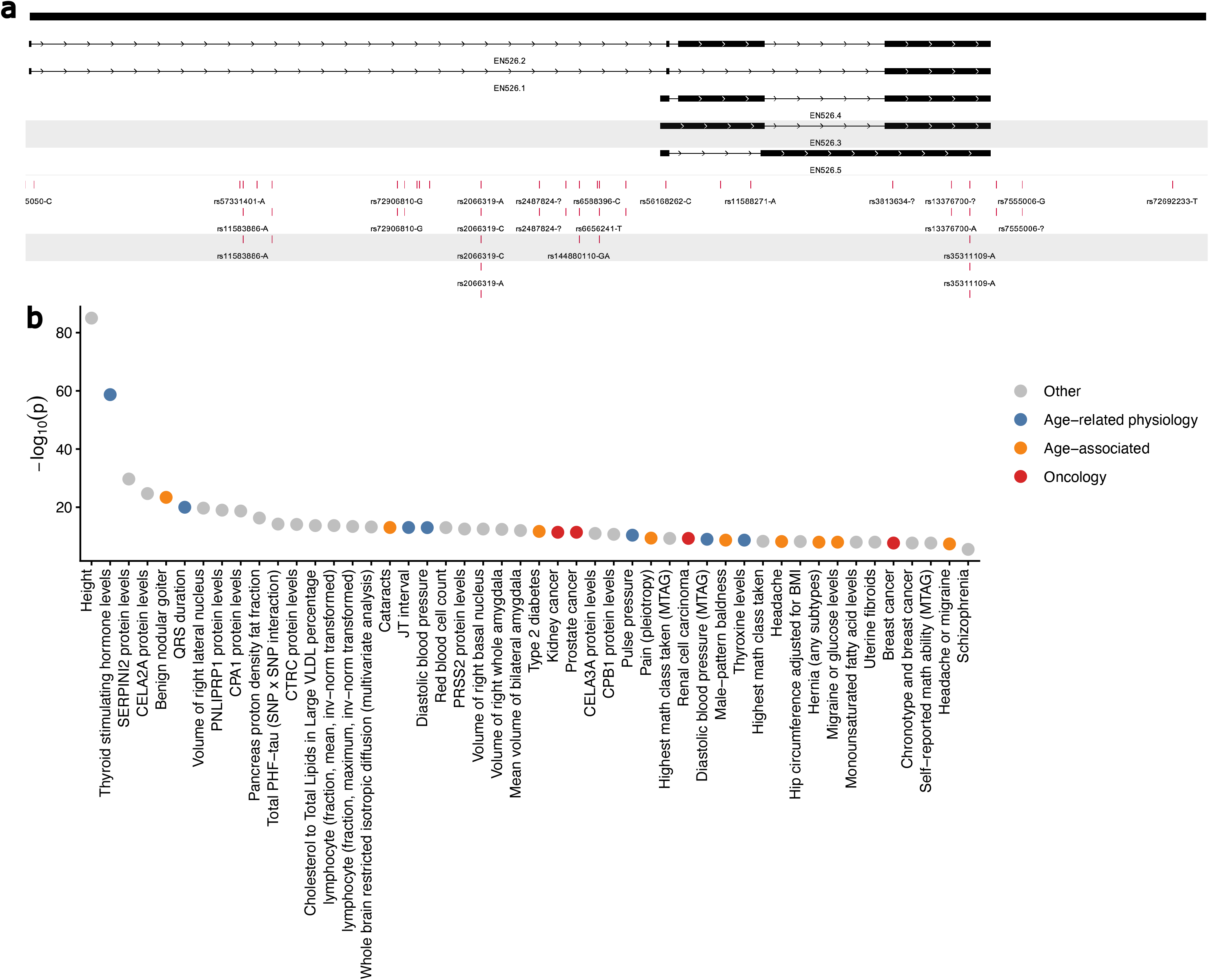
Genetic associations at the EN526 locus link to age-related traits and protein phenotypes. (**a**) Genomic view of the EN526 locus (hg38) showing EN526 transcript model and overlapping trait-associated variants identified from GWAS datasets. (**b**) Distribution of trait-associated variants overlapping the EN526 locus, plotted as −log10(p) for each associated trait. Traits are grouped and colour-coded by category, including age-related physiology, age-associated conditions, oncology, and other traits (Data: GWAS Catalog).

## 4. Discussion

Cellular senescence is a multifaceted stress response characterised by stable proliferative arrest, chromatin reorganisation, and widespread changes in gene expression and nuclear architecture ^66,67^. Despite extensive research, a unified mechanistic framework linking upstream inducers to downstream functional hallmarks remains elusive. In this study, we uncover a novel post-transcriptional regulatory axis involving eRNAs in senescence, anchored by a previously uncharacterised transcript, EN526, and its functional connection to stress responsive transcription factors, and CDKN2C (Fig. 1g, 2h). This EN526-CDKN2C axis integrates enhancer dynamics, RNA–RNA interactions, and downstream senescence phenotypes across molecular layers.

The role of eRNAs in transcriptional regulation is increasingly well established, with studies demonstrating enhancer-promoter looping, RNAPII recruitment, and local chromatin modification ^20,21,68–71^. However, emerging evidence suggests that some eRNAs are exported to the cytoplasm and may exert post-transcriptional effects ^22,23^, though few have been functionally characterised in this capacity. Here, we identify 21 SAeRs through integrated temporal and cross-context transcriptomic profiling. These eRNAs showed consistent dysregulation across senescence models, underscoring their potential generalisability. The minimal differential expression between early and late senescence (only 3 DE eRNAs, all upregulated) indicates that most eRNA remodelling has already occurred by early senescence, and late senescence largely maintains this transcriptional state. Their cytoplasmic localisation across cell types and RNA biotypes, especially within insoluble and membrane fractions, is supported by our recent findings that eRNAs are not solely nuclear-restricted and transcriptional regulators ^24^.

Among the SAeRs, EN526 emerged as a key node, exhibiting direct physical interactions with the mRNA of CDKN2C, an effector implicated in regulating cell cycle progression and apoptosis ^72,73^. CDKN2C is a cyclin-dependent kinase inhibitor that suppresses CDK4/6 activity, thereby enforcing RB1-dependent G1 arrest ^72^. Therefore, the downregulation of CDKN2C during senescence is consistent with the senescent cell state, which is defined by a stable proliferative arrest coupled with resistance to apoptosis. By dampening the expression of factors that actively promote either growth inhibition or cell death, senescent cells maintain a viable, yet non-dividing state, a hallmark of their long-term persistence in tissues. Crucially, these regulatory interactions were experimentally validated through crosslinking-based RNA–RNA interaction data (KARR-Seq, RIC-Seq, PARIS-Seq), curated in our eRNAkit resource ^23^. This structural evidence reinforces the idea that EN526 directly engages its mRNA targets to influence their stability and translation, providing a post-transcriptional mechanism that complements chromatin-level control. In the context of senescence, this post-transcriptional repression contributes to the reprogramming of gene networks that coordinate growth arrest with apoptotic resistance.

We traced repression of EN526 during senescence upstream of AT hook and bZIP TFs, a class of factors known to respond to cellular stress and implicated in apoptosis, inflammation, and chromatin regulation in senescence ^74–76^. We demonstrate that AT-hook and bZIP TFs associate with EN526 repression. Notably, this repression is not accompanied by decreased chromatin accessibility, indicating dynamic transcriptional repression rather than silencing via global chromatin compaction. The responses observed for BACH1 and SETBP1 despite the absence of detected binding at the EN526 locus suggest that candidate TFs may influence EN526 through different mechanisms. For example, bZIP TF activity depends on dimer composition ^77^, BACH1 depletion could therefore affect EN526 indirectly by altering the availability or activity of regulatory bZIP complexes. Alternatively, the absence of binding evidence may reflect cell-type-specific occupancy. Distinguishing direct occupancy from partner-mediated and indirect regulation will require targeted mechanistic investigation. Functionally, siRNA-mediated depletion of EN526 phenocopied senescence features across multiple axes: target gene repression (CDKN2C) and secretome changes. Importantly, these effects are mediated by exclusively depleting cytoplasmic EN526, confirming their post-transcriptional origin. This distinguishes our findings from prior enhancer studies which focus primarily on transcriptional activation or repression ^25,78^.

This work also addresses a major gap in the field, the lack of robust, mechanistically grounded senescence biomarkers. Recent studies have proposed multi-gene signatures ^79,80^, yet many rely on correlation rather than causality. Our proposed EN526-CDKN2C axis not only captures regulatory cause and consequence but also distinguishes senescence-specific responses from transient stress-related gene expression changes. This is particularly relevant in the context of aging, cancer therapy, and inflammatory disorders, where senescence and stress responses frequently overlap ^81^. Adding translational relevance, GWAS analyses revealed trait-associated SNPs in the EN526 enhancer locus, implicating it in red blood cell traits, glycaemic control, hormone regulation, and cardiovascular risk. These associations underscore the potential relevance of this regulatory region to human aging and disease and suggest that EN526 may act as a node integrating genetic susceptibility with downstream cellular phenotypes.

While these findings provide new insight into eRNA regulation of senescence, several limitations warrant consideration. First, our functional validation of EN526 effects focused on primary endothelial cells. Although the discovery analyses integrated datasets across multiple cell types and senescence contexts, the mechanistic perturbation experiments were conducted in a single cellular system. Future studies will be required to determine whether the EN526–CDKN2C regulatory axis operates similarly across other relevant cell types, including epithelial and immune lineages where senescence also plays prominent roles. Second, while multiple orthogonal datasets support physical interaction between EN526 and CDKN2C transcripts and our perturbation experiments demonstrate functional regulation, the precise molecular mechanism by which EN526 influences mRNA stability and translation remains to be fully elucidated. It will be important in future work to define the RNA structural features mediating this interaction and to determine whether additional RNA-binding proteins participate in this regulatory complex. Finally, the genetic associations identified within the EN526 enhancer locus derive from GWAS and phenome-wide datasets and therefore represent correlative links between genetic variation and aging-related traits. Although the overlap of trait-associated variants with the EN526 regulatory region and their linkage to CDKN2C support the physiological relevance of this pathway, functional studies will be required to determine whether these variants directly influence enhancer activity, EN526 expression, or downstream post-transcriptional regulation in age-related diseases.

## 5. Conclusion

In summary, this study identifies EN526 as a post-transcriptional effector of senescence and defines a novel, integrative axis: EN526-CDKN2C, that captures the senescence program across multiple levels of gene regulation. This axis deepens our mechanistic understanding of enhancer RNA biology in senescence and lays the foundation for functionally anchored biomarkers and targetable pathways in age-related disease.

## Ethical Statement

Not applicable

## CRediT authorship contribution statement

C.A.A. conceived and supervised the project and implemented the image analysis workflow. N.B, R.K and C.A.A curated the genomics datasets and performed the bioinformatic analyses. R.K, N.B, J.K, and C.A.A conducted the wet lab experiments. R.K, N.B, J.K, J.B, W.R and C.A.A analysed and interpreted the data. C.A.A wrote the manuscript. All authors edited and approved the final manuscript.

## Acknowledgements

Not applicable

## Data Availability

Publicly available datasets used in this study were obtained from the following repositories: ArrayExpress (E-MTAB-5403; https://www.ebi.ac.uk/arrayexpress/); Gene Expression Omnibus (GEO: GSE133292, GSE227766, GSE179810, GSE160356, GSE103938, GSE104666, GSE139963, GSE125218, GSE159539, GSE263439; https://www.ncbi.nlm.nih.gov/geo/); ENCODE (ENCSR226CVP and ENCSR109KMO; https://www.encodeproject.org/); ReMap2022 (https://remap.univ-amu.fr/); and the GWAS Catalog (https://www.ebi.ac.uk/gwas/). All data are available from the respective repositories under the accession numbers provided. Explicit usage information and full citations for all datasets are provided in the section “Integrative analysis of omics datasets across senescence contexts”. eRNA transcript models (GTF) and regulatory annotations for eRNA-mRNA interactions used in this study are available at https://github.com/AneneLab/eRNAkit. In addition, the script used for nuclear image segmentation and analysis is available at https://github.com/AneneLab/Eden “morphology”. This script has been optimised for typical images generated in our laboratory.

